# Genetic autonomy and low singlet oxygen yield support kleptoplast functionality in photosynthetic sea slugs

**DOI:** 10.1101/2021.02.02.429324

**Authors:** Vesa Havurinne, Maria Handrich, Mikko Antinluoma, Sergey Khorobrykh, Sven B. Gould, Esa Tyystjärvi

**Affiliations:** University of Turku, Department of Biochemistry / Molecular plant biology, Finland; Department of Biology, Heinrich-Heine-Universität, 40225 Düsseldorf, Germany

**Keywords:** Kleptoplasty, photoinhibition, photosynthetic sea slugs, PSII repair cycle, reactive oxygen species, singlet oxygen, *Vaucheria litorea*

## Abstract

*Elysia chlorotica* is a kleptoplastic sea slug that preys on *Vaucheria litorea*, stealing its plastids which then continue to photosynthesize for months inside the animal cells. We investigated the native properties of *V. litorea* plastids to understand how they withstand the rigors of photosynthesis in isolation. Transcription of specific genes in laboratory-isolated *V. litorea* plastids was monitored up to seven days. The involvement of plastid-encoded FtsH, a key plastid maintenance protease, in recovery from photoinhibition in *V. litorea* was estimated in cycloheximide-treated cells. *In vitro* comparison of *V. litorea* and spinach thylakoids was applied to investigate ROS formation in *V. litorea*. Isolating *V. litorea* plastids triggered upregulation of *ftsH* and translation elongation factor EF-Tu (*tufA*). Upregulation of FtsH was also evident in cycloheximide-treated cells during recovery from photoinhibition. Charge recombination in PSII of *V. litorea* was found to be fine-tuned to produce only small quantities of singlet oxygen (^1^O_2_). Our results support the view that the genetic characteristics of the plastids themselves are crucial in creating a photosynthetic sea slug. The plastid’s autonomous repair machinery is likely enhanced by low ^1^O_2_ production and by upregulation of FtsH in the plastids.

**Highlight:** Isolated *Vaucheria litorea* plastids exhibit upregulation of *tufA* and *ftsH*, key plastid maintenance genes, and produce only little singlet oxygen. These factors likely contribute to plastid longevity in kleptoplastic slugs.

## Introduction

Functional kleptoplasty in photosynthetic sea slugs depends on two major components: the first is a slug capable of stealing plastids and retaining them functional within its cells, the second a plastid with a specific genetic repertoire (de Vries *et al*., 2015). All species that are able to do this belong to the Sacoglossan clade (Rumpho *et al*., 2011; de Vries *et al*., 2014). These slugs are categorized based on their plastid retention times, i.e. no retention, short-term retention (hours to ~10 days) and long-term retention species (≥10 days to several months) (Händeler *et al*., 2009). The record holding slug *Elysia chlorotica* can retain plastids for roughly a year (Green *et al*., 2000). The mechanisms utilized by the slugs to selectively sequester plastids from their prey algae remain uncertain, although recent studies have shown that in *E. chlorotica* it is an active process reminiscent of that observed for symbiotic algae and corals (Chan *et al*., 2018). The slugs possibly rely on scavenger receptors and thrombospondin type-1 repeat proteins for plastid recognition (Clavijo *et al*., 2020).

The sacoglossan’s ability to sequester plastids tends to distract attention from the unique features of the sequestered organelle, forming the second component of a photosynthetic slug system. Long-term retention sea slugs are only able to maintain functional plastids from a restricted list of siphonaceous algae and usually from only one species. Some sacoglossa have a wide selection of prey algae, but long-term retention of plastids is still limited to specific algal sources (Christa *et al*., 2013; de Vries *et al*., 2013). The native robustness of some plastid types was noticed decades ago, and early on suggested to contribute to their functionality inside animals (Giles and Sarafis, 1972; Trench *et al*., 1973 *a*, b). Studies focusing on the specific properties of the algal plastids, however, are scarce. Reduction of the plastid genome (plastome) during evolution has stripped the organelle of many genes required for self-maintenance (Martin, 2003), but genomic analysis of algal plastomes suggests that three genes (*tufA, ftsH* and *psbA*) could be among those critical for plastid maintenance inside a slug cell (de Vries *et al*., 2013). Out of the three, *psbA* remains in all plastomes, including those of higher plants, whereas *tufA* and *ftsH* are encoded by most algal plastid genomes (Baldauf and Palmer, 1990; Oudot-Le Secq *et al*., 2007; de Vries *et al*., 2013). It has been suggested that the plastid-encoded translation elongation factor EF-Tu (*tufA*) helps maintain translation, specifically of the thylakoid maintenance protease FtsH (*ftsH*) involved in the repair cycle of Photosystem II (PSII) (de Vries *et al*., 2013). FtsH degrades the D1 protein (*psbA*) of damaged PSII before the insertion of *de novo* synthesized D1 into PSII (Mulo *et al*., 2012; Järvi *et al*., 2015). Without continuous replacement of the D1 protein, light-induced damage to PSII would rapidly curtail photosynthesis (Tyystjärvi and Aro, 1996).

Unlike all other known plastid sources of long-term retention slugs, *Vaucheria litorea* (Fig. 1), the sole prey of *E. chlorotica,* is not a green but a yellow-green alga, with plastids derived from red algal lineage through secondary endosymbiosis (Cruz *et al*., 2013) (Fig. 1B). The plastome of *V. litorea* possesses the three important genes (de Vries *et al*., 2013). Furthermore, the plastid-encoded FtsH of *V. litorea* has been shown to carry the critical metalloprotease domain that is not present in *ftsH* of other prey algae of long-term retention slugs (Christa *et al*., 2018). Here, we show that isolated plastids of *V. litorea* (Fig. 1C) maintain highly specific transcription of their genes, and exhibit adequate genetic autonomy in their capability to recover from light induced damage of PSII, i.e. photoinhibition. We also estimated reactive oxygen species (ROS) production in the thylakoid membranes of *V. litorea*. While our results highlight the importance of terminal electron acceptors downstream of Photosystem I (PSI) in limiting ROS production, we show that PSII of *V. litorea* is fine-tuned to decrease the yield of the highly reactive singlet oxygen (^1^O_2_). The consequences of our findings to light-induced damage and longevity of the plastids inside photosynthetic sea slugs are discussed in detail.

**Figure 1.**
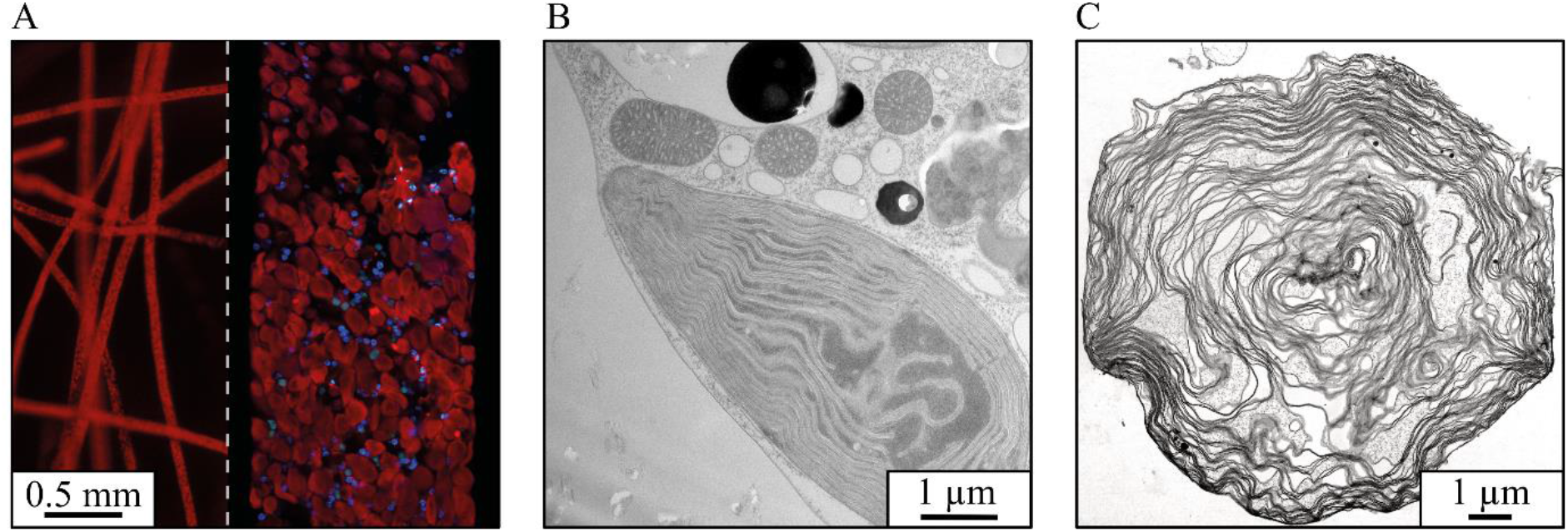
Microscope images of *V. litorea*, the main source of plastids for the photosynthetic sea slug *E. chlorotica*. (A) Chlorophyll autofluorescence (red) and nucleus specific dye fluorescence (blue) from *V. litorea* filaments, with a detail of a single filament on the right. (B) Transmission electron micrograph (TEM) showing a plastid *in vivo* in a *V. litorea* cell and in close proximity to several mitochondria, and in (C) an isolated single plastid.

## Materials and Methods

### Organisms and culture conditions

Spinach, *Spinacia oleracea* L. Matador (Nelson Garden, Tingsryd, Sweden), and *V. litorea* C. Agardh 1823 (SCCAP K-0379) were grown in SGC 120 growth chambers (Weiss Technik UK, Loughborough, United Kingdom) in 8/12 h and 12/12 h light/dark cycles, respectively. Growth light (Master TL-D 36W/840; Philips, Amsterdam, The Netherlands) photosynthetic photon flux density (PPFD) was set to 40 μmol m^−2^s^−1^ for both species. Temperature was maintained at 22 °C for spinach and 17 °C for *V. litorea.* Spinach plants used in the experiments were approximately 2 months old. *V. litorea* was grown in 500 ml flasks in f/2 culture medium (modified from Guillard and Ryther, 1962) made in 1% (m/v) artificial sea water (Sea Salt Classic, Tropic Marin, Wartenberg, Germany). *V. litorea* cultures were routinely refreshed by separating 1-4 g of inoculate into new flasks, and cultures used in the experiments were 1-2 weeks old. Nuclei of *V. litorea* were stained for microscopy with Hoechst 33342 (Thermo Scientific, Waltham, MA, USA) using standard protocols. *In vivo* transmission electron microscope (TEM) images were taken after freeze-etch fixation. The sea slug *Elysia timida* and its prey alga *Acetabularia acetabulum* were routinely maintained as described earlier (Schmitt *et al*., 2014; Havurinne and Tyystjärvi, 2020).

### Gene expression of isolated *V. litorea* plastids

Plastid isolation from *V. litorea* was performed based on Green *et al*. (2005). Briefly, filaments were cut to small pieces, resuspended in 40 ml of isolation buffer (see Table 1) and homogenized with ULTRA-TURRAX® (IKA, Staufen, Germany) using four short bursts at 8000 rpm. The homogenate was filtered twice through a layer of Miracloth (Calbiochem, Darmstadt, Germany), centrifuged (1900 x g, 5 min) and the pellet was resuspended in 1 ml of isolation buffer. Percoll solution containing 0.25 M sucrose was diluted to a 75 and 30% solution with 1xTE buffer containing 0.25 M sucrose. The sample was layered between the two dilutions and the assemblage was centrifuged (3500 x g, 20 min) in a swing-out rotor with no deceleration. Intact plastids were collected from the interphase and washed twice by centrifugation (2200 x g, 3 min) with isolation buffer lacking BSA. All steps were carried out at 4 °C in the dark. TEM imaging of the plastids was done after fixing the samples using glutaraldehyde and cryo-fixation followed by freeze substitution.

**Table 1.** Buffer solutions used in sample preparation and measurements.

| Identifier | Composition | Used in |
| --- | --- | --- |
| Plastid isolation buffer | 0.2 % BSA, 1 mM EDTA, 50mM Hepes-KOH pH 7.6, 1 mM MgCl<sub>2</sub>, 330 mM sorbitol | Plastid isolation, *in organello* gene expression |
| Thylakoid isolation buffer | 1 % BSA, 1 mM EDTA, 1 mM glycine betaine, 40 mM HEPES-KOH pH 7.4, 10 mM MgCl<sub>2</sub>, 0.3 M sorbitol | Thylakoid isolation |
| Osmotic shock buffer | 10 mM HEPES-KOH pH 7.4, 10 mM MgCl<sub>2</sub>, 5 mM sorbitol | Thylakoid isolation |
| Storage buffer | 10 mM HEPES-KOH pH 7.4, 10 mM MgCl<sub>2</sub>, 5 mM NaCl, 500 mM sorbitol | Thylakoid isolation, EPR |
| Photosystem measuring buffer | 1 M glycine betaine, 40 mM HEPES-KOH pH 7.4, 1 mM $\text{KH}_2\text{PO}_4$, 5 mM $\text{MgCl}_2$, 5 mM NaCl, 5 mM $\text{NH}_4\text{Cl}$, 330 mM sorbitol, | Thermoluminescence, flash oxygen evolution, fluorescence decay kinetics, $^1\text{O}_2$ production |
| Photoinhibition buffer | 1 M glycine betaine, 40 mM HEPES-KOH pH 7.4, 5 mM $\text{MgCl}_2$, 5 mM NaCl, 330 mM sorbitol | *In vitro* photoinhibition treatments, *in vitro* $\text{P}_{700}^+$ measurements |
| PSI measuring buffer | Photosystem measuring buffer + 0.3 mM 2,6-dichlorophenolindophenol (DCPIP), 0.01 mM 3-(3,4-dichlorophenyl)-1,1-dimethylurea (DCMU), 0.12 mM methyl viologen, 32 mM Na-ascorbate, 0.6 mM $\text{NaN}_3$ | Polarographic PSI activity measurements (oxygen consumption) |
| PSII measuring buffer | Photosystem measuring buffer + 0.5 mM 2,6-dichloro-1,4-benzoquinone (DCBQ), 0.5 mM hexacyanoferrate(III) | Polarographic PSII activity measurements (oxygen evolution) |

Plastids were kept in isolation buffer for seven days in routine culturing conditions. RNA was isolated at different time points using Spectrum™ Plant Total RNA Kit (Sigma-Aldrich, St. Louis, MO, USA). Aliquots with 50 ng RNA were subjected to DNAse treatment (Thermo Scientific), and treated aliquots amounting to 10 ng RNA were used for cDNA synthesis (iScript™ cDNA Synthesis Kit, BioRad, Hercules, CA, USA). Quantitative real-time PCR was carried out using a StepOnePlus (Applied Biosystems, Foster City, CA, USA) and reagents from BioRad. The primers used in the qPCR were designed using Primer3 (http://frodo.wi.mit.edu/primer3); the primer sequences are listed in Supplementary Table S1 at *JXB* online. Every reaction was done with technical triplicates and results were analyzed using the ΔΔCt method (Pfaffl, 2001), in which the qPCR data were double normalized to *rbcL* and time point 0 (immediately after plastid isolation).

### *In vivo* photoinhibition

The capacity to recover from photoinhibition was tested in spinach leaves and *V. litorea* cells in the presence of cycloheximide (CHI), a cytosolic translation inhibitor. Spinach leaf petioles were submerged in water containing 1 mM CHI and incubated for 24 h in the dark. The incubation was identical for *V. litorea* cells, except that the cells were fully submerged in f/2 medium supplemented with 1 mM CHI. Control samples were treated identically without CHI. The samples were then exposed to white light (PPFD 2000 μmol m^−2^ s^−1^) for 60 min and subsequently put to dark and thereafter low light (PPFD 10 μmol m^−2^ s^−1^) to recover for 250 min. Temperature was maintained at growth temperatures of both species using a combination of a thermostated surface and fans. The petioles of spinach leaves were submerged in water (-/+ CHI) during the experiments. Cell clusters of *V. litorea* were placed on top of the thermostated surface on a paper towel moistened thoroughly with f/2 medium (-/+ CHI). PSII activity was estimated by measuring the ratio of variable to maximum fluorescence (F_V_/F_M_) (Genty *et al*., 1989) with PAM-2000 (Walz, Effeltrich, Germany) fluorometer. During the light treatments, F_V_/F_M_ was measured from samples that were dark acclimated for <5 min, except for the final time point, where the samples were dark acclimated for 20 min. The light source used for all high-light treatments discussed in this study was an Artificial Sunlight Module (SLHolland, Breda, The Netherlands).

Membrane proteins were isolated at timepoints indicated in the figures. The same area where F_V_/F_M_ was measured (approximately 1 cm^2^) was cut out of the leaves/algal clusters and placed in a 1 ml Dounce tissue grinder (DWK Life Sciences, Millville, NJ, USA) filled with 0.5 ml of osmotic shock buffer (Table 1) and ground thoroughly. The homogenate was filtered through one layer of Miracloth and centrifuged (5000 x g, 5 min). The pellet containing the membrane protein fraction was resuspended in 50 μl of thylakoid storage buffer. The samples were stored at −80 °C until use. Membrane protein samples containing 1 μg total Chl were solubilized and separated by electrophoresis on a 10 % SDS-polyacrylamide gel using Next Gel solutions and buffers (VWR, Radnor, PA, USA). Proteins were transferred to Immobilon-P PVDF membranes (MilliporeSigma, Burlington, MA, USA). FtsH was immunodetected using antibodies raised against *Arabidopsis thaliana* FtsH5, reactive with highly homologous proteins FtsH1 and FtsH5, or FtsH2, reactive with FtsH2 and FtsH8 (Agrisera, Vännäs, Sweden). Western blots were imaged using goat anti-rabbit IgG (H+L) alkaline phosphatase conjugate (Life Technologies, Carlsbad, CA, USA) and CDP-star Chemiluminescence Reagent (Perkin-Elmer, Waltham, MA, USA). Protein bands were quantified with Fiji (Schindelin *et al*., 2012).

Experiments with *E. timida* were performed on freshly fed individuals. Slugs were kept in the dark overnight both in the absence and presence of 10 mg/ml lincomycin in 3.7 % artificial sea water and then exposed to high light (PPFD 2000 μmol m^−2^ s^−1^) in wells of a 24 well-plate filled with artificial sea water for 40 min. Temperature was maintained at 23 °C throughout the treatment. The slugs were then put to recover overnight in low light (PPFD <20 μmol m^−2^ s^−1^) in their growth conditions. F_V_/F_M_ was measured with PAM-2000 after a minimum 20 min dark period as described earlier (Havurinne and Tyystjärvi, 2020).

### Isolation of functional thylakoids for *in vitro* experiments

Functional thylakoids were isolated as described earlier (Hakala *et al*., 2005) after 24h dark incubation. One spinach leaf per isolation was ground in a mortar in thylakoid isolation buffer (Table 1). The homogenate was filtered through a layer of Miracloth and pelleted by centrifugation (5000 x g, 5 min). The pellet was resuspended in osmotic shock buffer, centrifuged (5000 x g, 5 min) and the resulting pellet was resuspended in thylakoid storage buffer. Chl concentration was determined spectrophotometrically in 90 % acetone using extinction coefficients for Chls *a* and *b* (Jeffrey and Humphrey, 1975). Thylakoid isolation from *V. litorea* was performed using the same procedure, by grinding 2-5 g of fresh cell mass per isolation. The cell mass was briefly dried between paper towels before grinding. Chl concentration from *V. litorea* thylakoids was determined in 90% acetone using coefficients for Chls *a* and *c1* + *c2* (Jeffrey and Humphrey, 1975). Protein concentrations of the thylakoid suspensions were determined with DC™ Protein Assay (Bio-Rad, Hercules, CA, USA). Thylakoids used in functional experiments were kept on ice in the dark and always used within a few hours of isolation.

### Photosystem stoichiometry

Photosystem stoichiometry was measured from thylakoid membranes with an EPR spectroscope Miniscope MS5000 (Magnettech GmbH, Berlin, Germany) as described earlier (Tiwari *et al*., 2016; Nikkanen *et al*., 2019). EPR spectra originating from oxidized tyrosine-D residue of PSII (Tyr_D_^+^) and reaction center Chl of PSI (P_700_^+^) of concentrated thylakoid samples (2000 μg Chl ml^−1^ in storage buffer) were measured in a magnetic field ranging from 328.96 to 343.96 mT during illumination (PPFD 4000 μmol m^−2^ s^−1^) (Lightningcure LC8; Hamamatsu Photonics, Hamamatsu City, Japan) and after a subsequent 5 min dark period in the absence and presence of 50 μM 3-(3,4-dichlorophenyl)-1,1-dimethylurea (DCMU). The dark stable Tyr_D_^+^ EPR signal (PSII signal), measured after the post illumination period in the absence of DCMU, and the P_700_^+^(PSI signal), measured during illumination in the presence of DCMU, were double integrated to determine photosystem stoichiometry.

### *In vitro* photoinhibition

For *in vitro* photoinhibition experiments, thylakoids were diluted to a total Chl concentration of 100 μg ml^−1^ in photoinhibition buffer (Table 1), and 1 ml sample was loaded into a glass beaker submerged in a water bath kept at 22 °C. The samples were exposed to white light (PPFD 1000 μmol m^−2^ s^−1^) and mixed with a magnet during the 60 min treatments. Aliquots were taken at set intervals to determine PSI or PSII activities using a Clark-type oxygen electrode (Hansatech Instruments, King’s Lynn, England). The sample concentration in the activity measurements was 20 μg total Chl ml^−1^ in 0.5 ml of PSI or PSII measuring buffer (Table 1). PSI activity was measured as oxygen consumption, whereas PSII activity was measured as oxygen evolution. Both activities were measured at 22 °C in strong light (PPFD 3200 μmol m^−2^ s^−1^) from a slide projector. The rate constant of PSII photoinhibition (k_PI_) was obtained by fitting the loss of oxygen evolution to a first-order reaction equation with Sigmaplot 13.0 (Systat Software, San Jose, CA, USA), followed by dark correction, i.e. subtraction of the dark inactivation rate constant from the initial k_PI_.

Lipid peroxidation was measured by detecting malondialdehyde (MDA) formation (Heath and Packer, 1968). A thylakoid suspension aliquot of 0.4 ml was mixed with 1 ml of 20 % trichloroacetic acid containing 0.5 % thiobarbituric acid, incubated at 80 °C for 30 min and cooled down on ice for 5 min. Excess precipitate was pelleted by centrifugation (13500 x g, 5 min), and the difference in absorbance between 532 and 600 nm (Abs_532-600_) was measured as an indicator of the relative amount of MDA in the samples. Protein oxidation was determined by detecting protein carbonylation with Oxyblot™ Protein Oxidation Detection Kit (MilliporeSigma, Burlington, MA, USA). Thylakoid aliquot amounting to a protein content of 45 μg was taken at set time points and 10 mM dithiothreitol was used to prevent further protein carbonylation. The samples were prepared according to the manufacturer’s instructions and proteins were separated in 10 % Next Gel SDS-PAGE (VWR). Carbonylated proteins were detected with Immobilon Western Chemiluminescent HRP Substrate (MilliporeSigma).

The maximum oxidation of P_700_ (P_M_) was estimated in an additional experiment. Thylakoids equivalent to 25 ug Chl in 50 μl of photoinhibition buffer were pipetted on a Whatman filter paper (grade 597; Cytiva, Marlborough, MA, USA). The filter was placed inside the lid of a plastic Petri dish, and the bottom of the Petri dish was placed on top of the lid. Photoinhibition buffer was added to the sample from the small openings on the sides of the assemblage. The thylakoids were then illuminated with high light (PPFD 1000 μmol m^−2^ s^−1^) and the temperature was maintained at 22 °C using a thermostated surface. F_V_/F_M_ and P_M_ were measured using a 700 ms high-light pulse (PPFD 10000 μmol m^−2^ s^−1^) with Dual-PAM 100 (Walz) (Schreiber, 1986; Schreiber and Klughammer, 2008) at set intervals. The high-light treated samples were dark acclimated for <5 min prior to the measurements.

### ^1^O_2_ measurements

^1^O_2_ was measured from thylakoids diluted to 100 μg total Chl ml^−1^ in 0.3 ml of photosystem measuring buffer, using the histidine method described earlier (Telfer *et al*., 1994; Rehman *et al*., 2013). Continuously stirred thylakoid samples were exposed to high light (PPFD 3200 μmol m^−2^ s^−1^) from a slide projector at 22 °C in the presence and absence of 20 mM histidine. Oxygen consumption was measured for 60 s using an oxygen electrode (Hansatech), and the difference in the oxygen consumption rates in the presence and absence of histidine was taken as an indicator of ^1^O_2_ production. PSII electron transfer activity (H_2_O to DCBQ) in the same conditions was 124.7 (SE±15.4) and 128.4 (SE±10.7) μmol O_2_ mg Chl^−1^ h^−1^ in spinach and *V. litorea* samples, respectively, containing 20 μg Chl ml^−1^.

### PSII charge recombination measurements

Flash-induced oxygen evolution was recorded at room temperature using a Joliot-type bare platinum oxygen electrode (PSI, Brno, Czech Republic) (Joliot and Joliot, 1968) from thylakoids diluted in photosystem measuring buffer to 50 μg Chl ml^−1^ and supplemented with 50 mM KCl, essentially as described in Antal *et al*. (2009). 200 μl of sample was pipetted on the electrode and kept in the dark for 10 min before the measurements. The samples were then exposed to a flash train consisting of 15 single-turnover flashes (4 ns/pulse) at one second intervals, provided by a 532 nm Nd:YAG laser (Minilite, Continuum, San Jose, CA, USA). Charge recombination within PSII was probed by exposing the samples to a preflash and different dark times between the preflash and the flash train used for recording the oxygen traces.

The decay of Chl *a* fluorescence yield after a 30 μs single turnover flash (maximum PPFD 100 000 μmol m^−2^ s^−1^) were measured at room temperature from 1 ml samples of thylakoids using FL200/PS fluorometer (PSI). Measurement length was 120 s and 8 datapoints/decade were recorded (2 in the presence of DCMU). The first datapoint was recorded 150 μs after the flash. Single turnover flash and measuring beam voltages were set to 100 % and 60 % of the maximum, respectively. The samples were diluted in photosystem measuring buffer to a total Chl concentration of 20 μg ml^−1^. A set of samples was poisoned with 20 μM DCMU to block electron transfer at the reducing side of PSII.

Thermoluminescence was measured from thylakoids using a custom setup (Tyystjärvi *et al*., 2009). Thylakoids were diluted to a total Chl concentration of 100 μg ml^−1^ in photosystem measuring buffer (Table 1) in the presence and absence of 20 μM DCMU, and a volume of 100 μl was pipetted on a filter paper disk that was placed inside the cuvette of the measuring apparatus. The samples were dark acclimated for 5 min before the onset of cooling to −20 °C by a Peltier element (TB-127-1,0-0,8; Kryotherm, Carson City, NV, USA). The samples were then exposed to a flash (E = 1 J) from a FX-200 Xenon lamp (EGandG, Gaithersburg, MD, USA) and heated at a rate of 0.47 °C s^−1^ up to 60 °C while simultaneously recording luminescence emission.

### *In vivo* P_700_ redox kinetics

Redox kinetics of P_700_ were measured as described by Shimakava *et al*., (2019) using Dual-PAM 100 (Walz). Spinach plants and *V. litorea* cells were kept in darkness for at least 2 h before the measurements. Anaerobic conditions were obtained using a custom cuvette described in Havurinne and Tyystjärvi (2020). For spinach leaf cutouts, the cuvette was flushed with nitrogen. A combination of glucose oxidase (8 units/ml), glucose (6 mM) and catalase (800 units/ml) in f/2 culture medium was used to create anaerobic conditions for *V. litorea* cells. All samples were treated with 15 s of far red light (PFD 120 μmol m^−2^ s^−1^) and a subsequent darkness lasting 25 s prior to firing a high-light pulse (780 ms, PPFD 10 000 μmol m^-2^ s^-1^).

## Results

### Isolated *V. litorea* plastids maintain regulated gene expression

Laboratory-isolated *V. litorea* plastids exhibited differentially regulated gene expression even after seven days in isolation (Fig. 2). The orientations of selected genes in *V. litorea* plastome are shown in Fig. 2A. PSII core subunit genes *psbA, psbB*, *psbC* and *psbD* were downregulated after day 3 of the isolation period, while *psbB* and *psbD*, encoding CP47 and D2 proteins of PSII, reached a stationary level after five days, and the transcription of the genes encoding PSII proteins CP43 (*psbC*) and D1 (*psbA*) were among those downregulated most significantly (Fig. 2B). The main protein of PSII targeted for degradation after photoinhibition is D1, whereas release of CP43 from the PSII core has been suggested to precede D1 degradation in higher plants (Aro *et al*., 2005). One gene, *psbH*, encoding a small PSII subunit involved in proper PSII assembly in cyanobacteria (Komenda *et al*., 2005), exhibited stationary transcript levels throughout the isolation period, similar to the gene encoding PSI reaction center subunit PsaA. Transcription of *ftsH* and *tufA*, encoding the maintenance protease FtsH and the translation elongation factor EF-Tu, followed an upward trajectory throughout the experiment (Fig. 2B). We also tested the genetic autonomy of plastids sequestered by *E. timida* that feeds on *A. acetabulum*. Subjecting the slugs to high light for 40 min resulted in a drastic decrease in PSII photochemistry (F_V_/F_M_), but the kleptoplasts inside the slugs were capable of restoring PSII activity back to 78 % of the initial level during a 20 h recovery period. Subjecting the slugs to lincomycin, a plastid specific translation inhibitor (Mulo *et al*., 2003), however, almost completely prevented the recovery (Fig. 2C).

**Figure 2.**
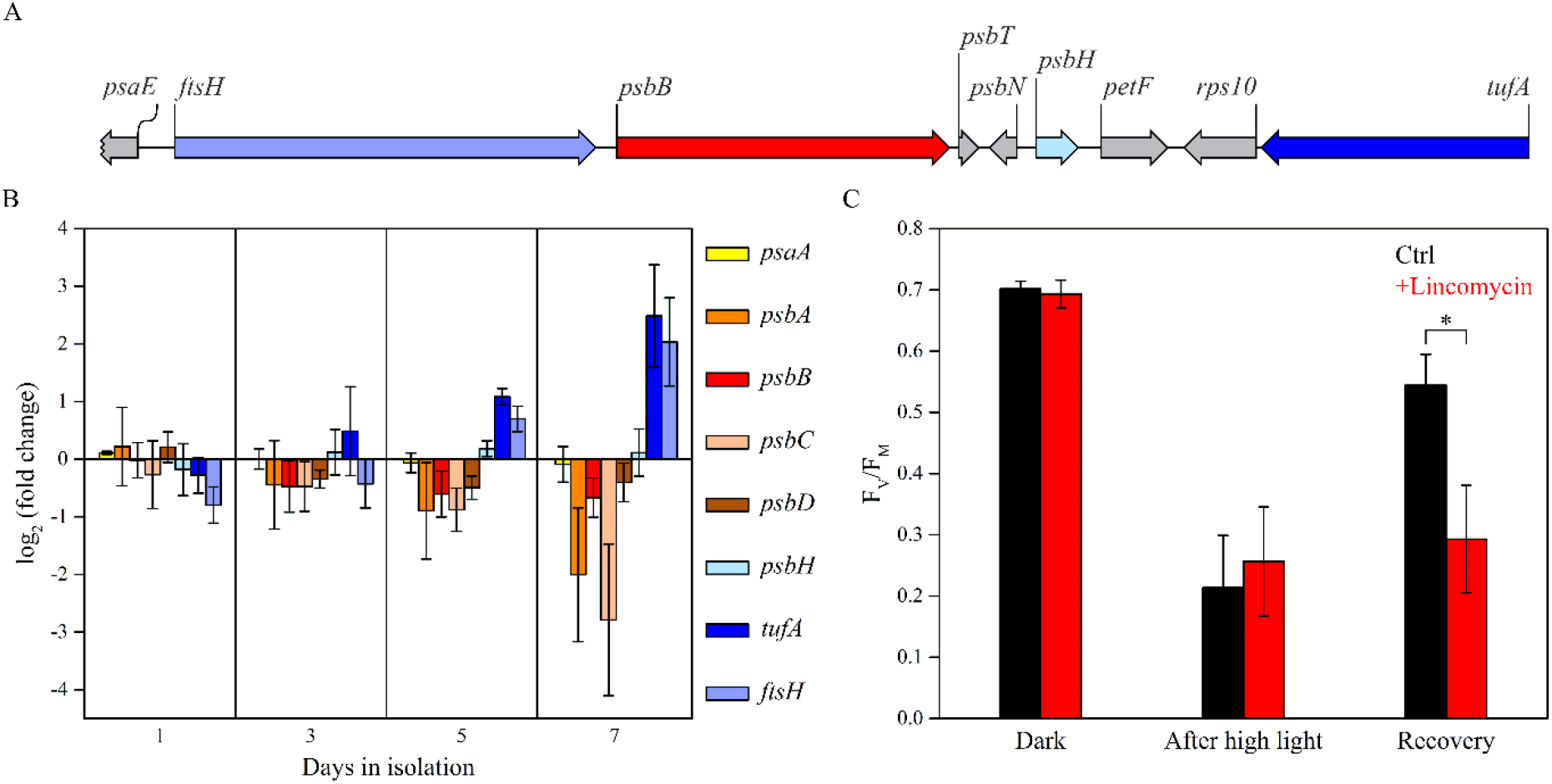
Transcription of plastid encoded genes in isolated *V. litorea* plastids and the autonomy of kleptoplasts inside the sea slug *E. timida*. (A) Orientation of specific genes inspected in (B) in *V. litorea* plastid genome. (B) Amounts of transcripts of selected genes during a period of seven days in isolation buffer; each transcript has been compared to the amount measured immediately after plastid isolation. (C) Maximum quantum yield of PSII photochemistry (F_V_/F_M_) measured at different timepoints of the photoinhibition treatment (40 min, PPFD 2000 μmol m^−2^s^−1^) and after overnight recovery (PPFD< 20 μmol m^−2^ s^−1^) in *E. timida* slug individuals in the absence and presence of lincomycin. The data in panels (B) and (C) are averages from three and four biological replicates, respectively. Error bars indicate standard deviation. An asterisk indicates a statistically significant difference between the two groups (Welch’s t-test, P<0.005).

### FtsH translation is enhanced in functionally isolated plastids of *V. litorea* during recovery from photoinhibition

Treating spinach leaves with CHI, a cytosolic translation inhibitor, resulted in faster loss of PSII activity in high light (Fig. 3A). Also PSII repair was impaired by CHI in spinach. *V. litorea* showed almost no effect of CHI during the same photoinhibition and recovery treatment (Fig. 3B). Using two different FtsH antibodies (FtsH 1+5 and FtsH 2+8), we tested the possible involvement of plastid-encoded FtsH of *V. litorea* in the unaffected PSII photochemistry in CHI treated samples. There were no differences in the relative protein levels of FtsH between control and CHI treated spinach during the experiment (Fig. 3C). Genes for FtsH reside in the nucleus in spinach, and our results suggest that the CHI treatment did not inhibit cytosolic translation in the leaves entirely, although *de novo* synthesis of proteins could not be tested by radiolabeling experiments. In *V. litorea,* CHI treatment increased FtsH levels towards the end of the experiment (Fig. 3D). This suggests that not only is expression of plastome genes active in functionally isolated plastids of *V. litorea*, but the translation of specific genes such as *ftsH* can be upregulated when the plastids are deprived from normal cytosolic governance.

**Figure 3.**
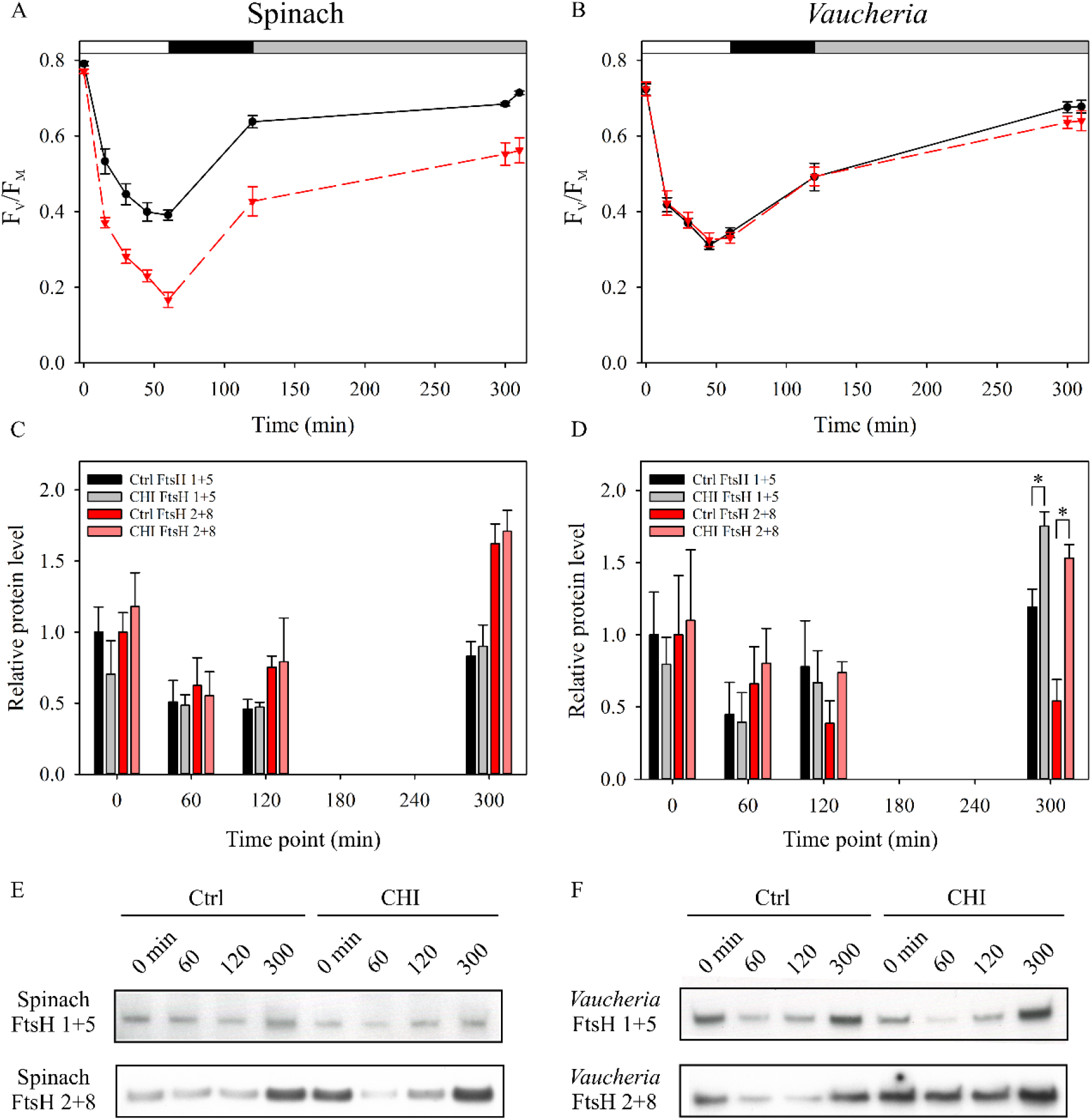
*V. litorea* recovers from photoinhibition of PSII in the presence of cycloheximide, a cytosolic translation inhibitor, and exhibits upregulation of FtsH. Quantum yield of PSII photochemistry (F_V_/F_M_) during photoinhibition treatment and subsequent recovery of (A) spinach and (B) *V. litorea* in the absence (ctrl; black) and presence of CHI (red). 0 min timepoint was measured before the onset of high-light treatment, 60 min timepoint after the high-light treatment (PPFD 2000 μmol m^−2^s^−1^), 120 min timepoint after subsequent dark recovery, and 300 min timepoint after recovery in dim light (10 μmol m^−2^s^−1^). The final timepoint at 310 min was measured after additional 10 min dark acclimation. The white, black and gray bars on top indicate the high-light treatment, dark and dim light periods, respectively. (C) Relative levels of FtsH in spinach and (D) *V. litorea* during the experiment, as probed by antibodies raised against *A. thaliana* FtsH 5 (FtsH 1+5; black and grey bars for ctrl and CHI treatments, respectively) and FtsH 2 (FtsH 2+8; red and light red bars for ctrl and CHI treatments). The light treatment regime up to 300 min was the same as in (A) and (B). Significant differences between treatments are indicated by an asterisk (Welch’s t-test, P<0.05, n=3). (E, F) Representative FtsH Western blots from spinach and *V. litorea,* respectively, used for protein quantification in panels (C) and (D). All data in (A) to (D) represent averages from at least three independent biological replicates and the error bars represent SE.

### Thylakoids of *V. litorea* exhibit moderate photoinhibition of PSII and elevated ROS damage, but produce little ^1^O_2_

Basic photosynthetic parameters of isolated thylakoids from spinach and *V. litorea* are shown in Table 2. Photoinhibition of PSII during a 60 min high-light treatment of isolated thylakoids proceeded according to first-order reaction kinetics (Tyystjärvi and Aro, 1996) in both species (Fig. 4A). However, spinach thylakoids were more susceptible to damage, as indicated by the larger rate constant of dark-corrected PSII photoinhibition (k_PI_) (Table 2). General oxidative stress assays of lipids and proteins of the thylakoid membranes exposed to high light showed more ROS damage in *V. litorea* than in spinach thylakoids during the treatment (Fig. 4B,C). Measurements of ^1^O_2_ production, the main ROS produced by PSII (Krieger-Liszkay, 2005; Pospíšil, 2012), from isolated thylakoids showed that the rate of ^1^O_2_ production in *V. litorea* is only half of that witnessed for spinach (Fig. 5A). This suggests that the main ROS, causing the *in vitro* oxidative damage to lipids and proteins (Fig. 4B,C) in *V. litorea*, are partially reduced oxygen species produced by PSI.

**Table 2.** Photosynthesis-related parameters of isolated spinach and *V. litorea* thylakoid membranes. The EPR spectra used for estimating the PSI/PSII ratio are shown in Supplementary Fig. S1. The indicated PSII and PSI activities are averages from all initial activity measurements of untreated control samples discussed in this publication. The k_PI_ value was determined from first-order reaction fits of the photoinhibition data in Fig. 4A, and corrected by subtracting the first-order rate constant of PSII inhibition in the dark (Supplementary Fig. S2). All values are averages from a minimum of three biological replicates and SE is indicated in parentheses.

| Organism | PSI/PSII | PSII activity<br>(H<sub>2</sub>O to | PSI activity<br>(DCPIP to methyl | $k_{PI}$ (min<sup>-1</sup>) |
| --- | --- | --- | --- | --- |

| | | DCBQ;<br>$\mu\text{mol O}_2$<br>evolved mg<br>$\text{Chl}^{-1} \text{h}^{-1}$) | viologen;<br>$\mu\text{mol O}_2$<br>consumed mg<br>$\text{Chl}^{-1} \text{h}^{-1}$) | |
| --- | --- | --- | --- | --- |
| Spinach | 2.438<br>($\pm 0.100$) | 200.12<br>($\pm 11.53$) | 758.22<br>($\pm 77.14$) | 0.0289<br>($\pm 0.002$) |
| *V. litorea* | 2.343<br>($\pm 0.090$) | 244.54<br>($\pm 15.71$) | 797.36<br>($\pm 93.73$) | 0.0148<br>($\pm 0.001$) |

**Figure 4.**
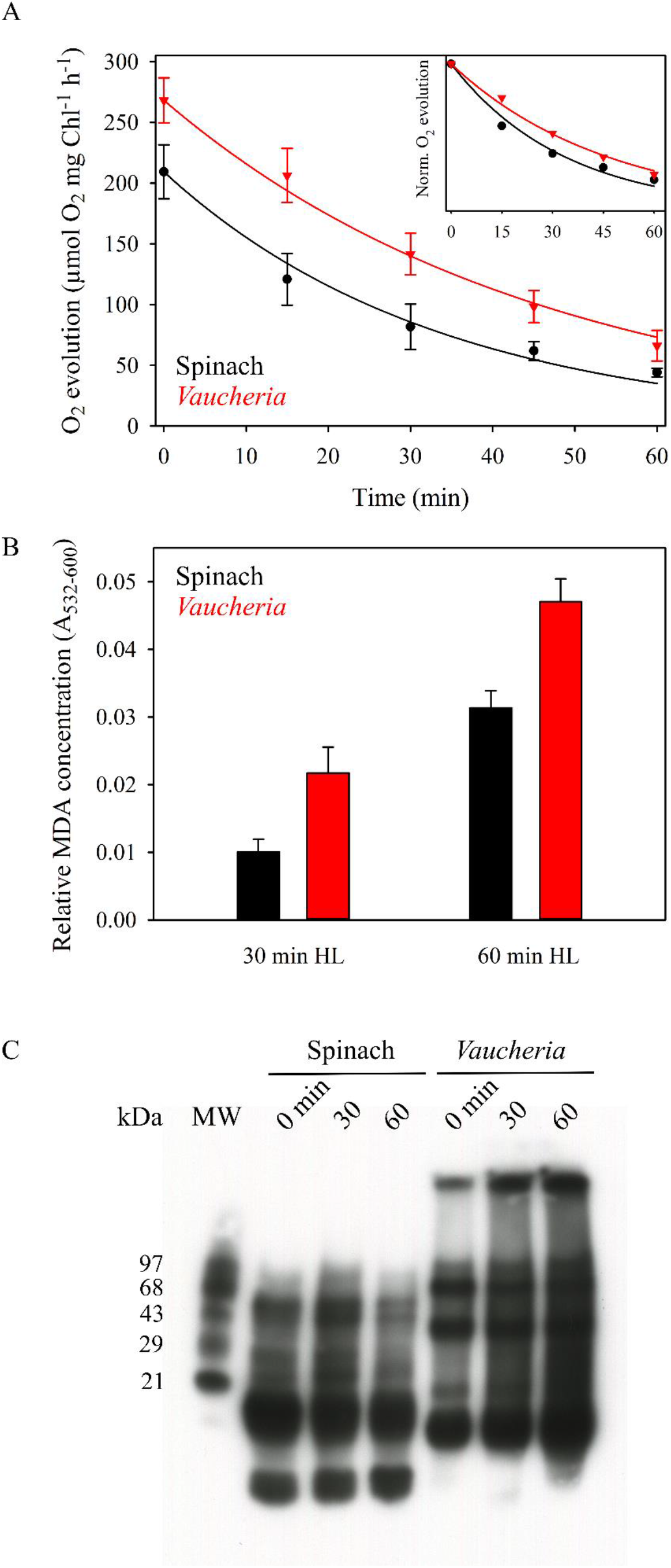
*In vitro* photoinhibition of PSII and ROS production in spinach (black) and *V. litorea* (red) thylakoids in high light. (A) Photoinhibition of PSII in high light (PPFD 1000 μmol m^−2^s^−1^), as estimated by oxygen evolution. The curves show the best fit to a first order reaction in spinach and *V. litorea*. Data normalized to the initial oxygen evolution rates are shown in the inset to facilitate comparison. Dark control experiments, shown in supplementary Fig. S2, indicated a 4.9 % (SE±3.6, n=3) and 27.5 % (SE±6.7, n=3) loss of PSII activity after 60 min in the dark for spinach and *V. litorea*, respectively. (B) Lipid peroxidation after 30 and 60 min of high-light treatment in spinach and *V. litorea*, as indicated by MDA formation. MDA formed during dark control treatments were subtracted from the high-light treatment data. (C) A representative Oxyblot™ assay of protein carbonylation during the high-light treatment. Each data point in panels (A, B) represents an average from a minimum of three biological replicates and the error bars indicate SE.

**Figure 5.**
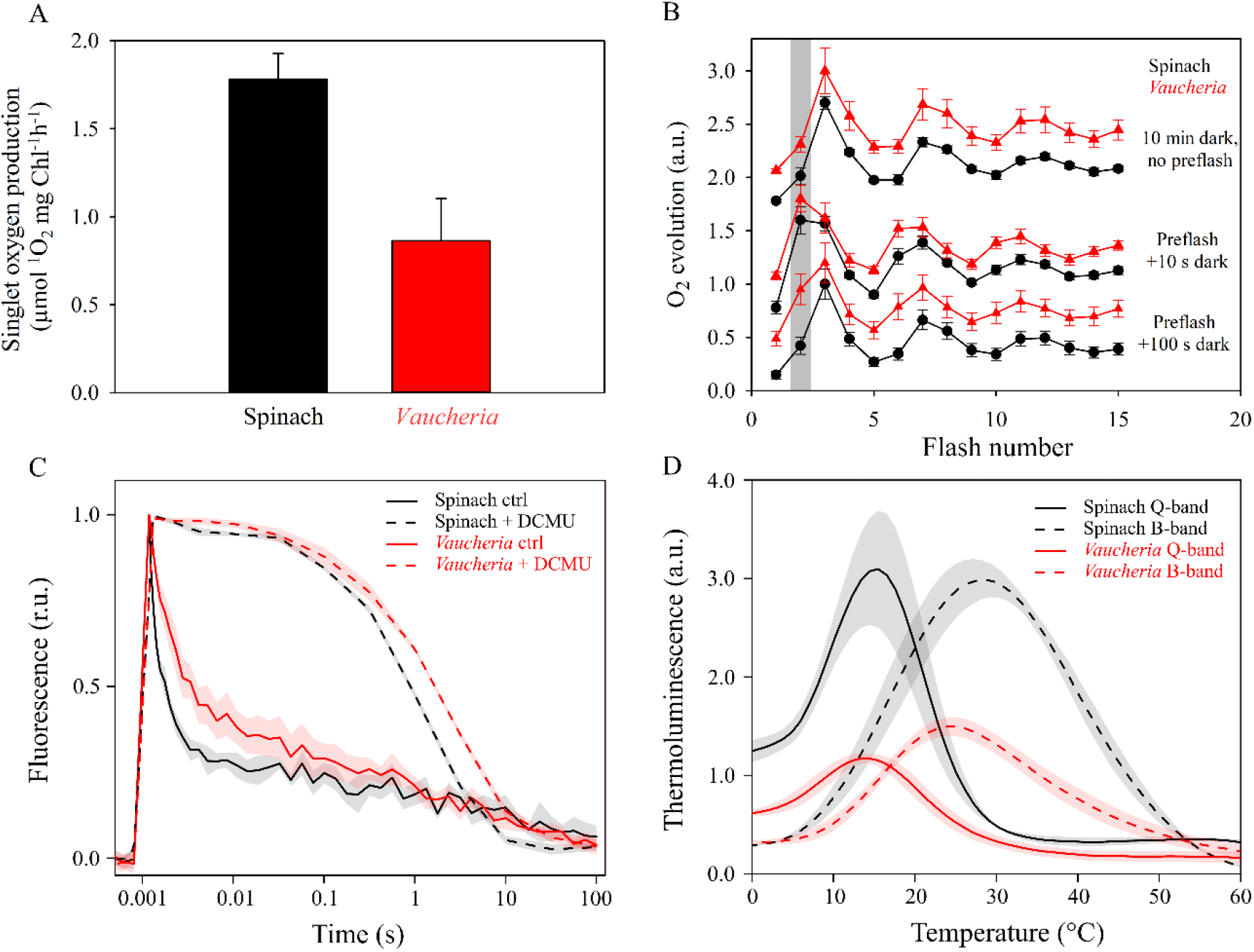
*V. litorea* thylakoids produce little ^1^O_2_ and exhibit slow charge recombination of PSII. (A) ^1^O_2_ production in spinach (black) and *V. litorea* (red) thylakoid membranes. (B) Flash oxygen evolution after different preflash treatments in spinach and *V. litorea* thylakoids. The grey bar highlights the oxygen yield instigated by the second flash, an indicator of charge recombination reactions taking place during the dark period between a preflash and the measuring flash sequence. Oxygen traces were double normalized to the first (zero level) and third flash and shifted in Y-axis direction for clarity. (C) Chl fluorescence decay kinetics after a single turnover light pulse in untreated (solid lines) and DCMU poisoned (dashed lines) thylakoids, double normalized to zero level before the onset of the pulse and maximum fluorescence measured 150 μs after the pulse. (D) Q (solid lines) and B band (dashed lines) of thermoluminescence, measured in the presence and absence of DCMU, respectively. All data in (A-C) represent averages from at least three biological replicates. Thermoluminescence data in (D) are from three replicates obtained from pooled thylakoid batches isolated from three plants/algae flasks. Error bars and shaded areas around the curves show SE.

### *V. litorea* produces only little ^1^O_2_, likely due to slow PSII charge recombination

We probed charge recombination reactions within PSII using three different methods to investigate the role of PSII in the low ^1^O_2_ yield in *V. litorea* thylakoids (Fig. 5A). First, we measured flash-induced oxygen evolution from isolated thylakoids of spinach and *V. litorea*. After 10 min dark acclimation, thylakoids from both species exhibited a typical pattern of oxygen evolution, i.e. the third flash caused the highest oxygen yield due to the predominance of the dark-stable S_1_ state of the oxygen evolving complex (OEC), after which the oxygen yield oscillated with a period of four until dampening due to misses and charge recombination reactions (Fig. 5B, top curves). A single turnover pre-flash treatment makes S_2_ the predominant state. A 10 s dark period after the pre-flash treatment was not long enough to cause noticeable changes in the S-state distribution in either species, as can be seen from the middle curves of Fig. 5B, where the second flash of the flash train causes the highest yield of oxygen. In spinach, 100 s darkness after the pre-flash treatment resulted in nearly complete restoration of the original S-states, whereas in *V. litorea* the second flash still yielded a considerable amount of oxygen (Fig. 5B, bottom curves). This is likely due to slow charge recombination between Q_B_^−^ and the S_2_ state of the OEC in *V. litorea* (Pham *et al*., 2019). The modeled percentage S-state distributions of OEC from spinach and *V. litorea* after different dark times between the pre-flash and the flash train are shown in Supplementary table S2.

Next, we measured the decay of Chl *a* fluorescence yield after a single turnover flash from thylakoids in the absence and presence of the PSII electron transfer inhibitor DCMU. Fluorescence decay in the absence of DCMU reflects Q_A_^−^ reoxidation mainly by electron donation to Q_B_ and Q_B_^−^. In the presence of DCMU, fluorescence decay is indicative of Q_A_^−^ reoxidation through various charge recombination reactions (Mamedov *et al*., 2000), some of which generate the harmful triplet P_680_ Chl through the intermediate P_680_+Pheo^−^ radical pair (Sane *et al*., 2012). The decay of fluorescence yield was slower in *V. litorea* thylakoids than in spinach both in the absence and presence of DCMU (Fig. 5C). In the absence of DCMU, the slower kinetics in *V. litorea* shows that electron transfer from Q_A_^−^ to Q_B_ is not as favorable as in spinach. The slow decay of fluorescence in the presence of DCMU indicates slow S_2_Q_A_^−^ charge recombination.

Thermoluminescence Q and B bands from thylakoids in the presence and absence of DCMU, respectively, were also measured. For a description on the interpretation of thermoluminescence data, see Tyystjärvi and Vass, (2004) and Sane *et al*., (2012). Briefly, the thylakoid samples were dark acclimated for 5 min, cooled down to −20 °C, flashed with a single turnover Xenon flash and then heated with a constant rate. The luminescence emitted by the samples at different temperatures is proportional to the rate of the luminescence-producing charge recombination reactions between the S-states of the OEC and downstream electron acceptors, more specifically S_2_/Q_A_^−^ (Q band) and S_2,3_/Q_B_^−^ (B band). The Q and B band emission peaks in spinach were at 15 and 28 °C, whereas in *V. litorea* they were at 14 and 24 °C (Fig. 5D). The lower peak temperatures in *V. litorea* would actually suggest that both Q_A_^−^ and Q_B_^−^ are less stable at room temperature in *V. litorea* than in spinach. However, the multiple pathways of recombination (Rappaport and Lavergne, 2009) obviously allow the luminescence-producing minor pathway to suggest destabilization of Q_A_^−^ in *V. litorea* (Fig. 5D) even if the total recombination reaction is slower in *V. litorea* than in spinach (Fig. 5B,C and Supplementary table S2). The thermoluminescence signal intensity was lower in *V. litorea* than in spinach, suggesting that the luminescence-producing reaction has a low yield in *V. litorea*. The narrow energy gap between Q_A_ and Q_B_ in *V. litorea* favors the probability of an electron residing with QA. Furthermore, a small Q_A_−Q_B_ energy gap also increases the probability that S_3_Q_B_^−^ or S_2_Q_B_^−^ recombine directly and non-radiatively without producing triplet P_680_ and subsequently ^1^O_2_ (Ivanov *et al*., 2003; Sane *et al*., 2003; Ivanov *et al*., 2008; Sane *et al*., 2012).

### *In vitro* high-light treatment lowers electron donation to methyl viologen and maximal oxidation of P_700_ in *V. litorea*

When PSI activity was estimated as electron transfer from DCPIP to methyl viologen (oxygen consumption), spinach PSI remained undamaged during *in vitro* high-light treatment, while *V. litorea* seemed highly susceptible to photoinhibition of PSI (Fig. 6A,B). We repeated the photoinhibition experiment, but this time PSII and PSI activities were monitored with Chl fluorescence and P_700_ absorption changes. Again, thylakoid membranes of spinach were more sensitive to photoinhibition of PSII during the high-light treatment than *V. litorea* (Fig. 6C,D). However, this time PSI functionality of both species decreased similarly when estimated as the maximum oxidation of P_700_ (P_M_). The decrease in PM was strong during the first 15 (*V. litorea*) or 30 min (spinach) of the light treatment, whereafter P_M_ remained at a somewhat stationary level (Fig. 6C,D). The decrease in P_M_ depended on electron transfer from PSII, as P_M_ did not decrease in high light in spinach thylakoids in the presence of DCMU (Supplementary Fig. S4).

**Figure 6.**
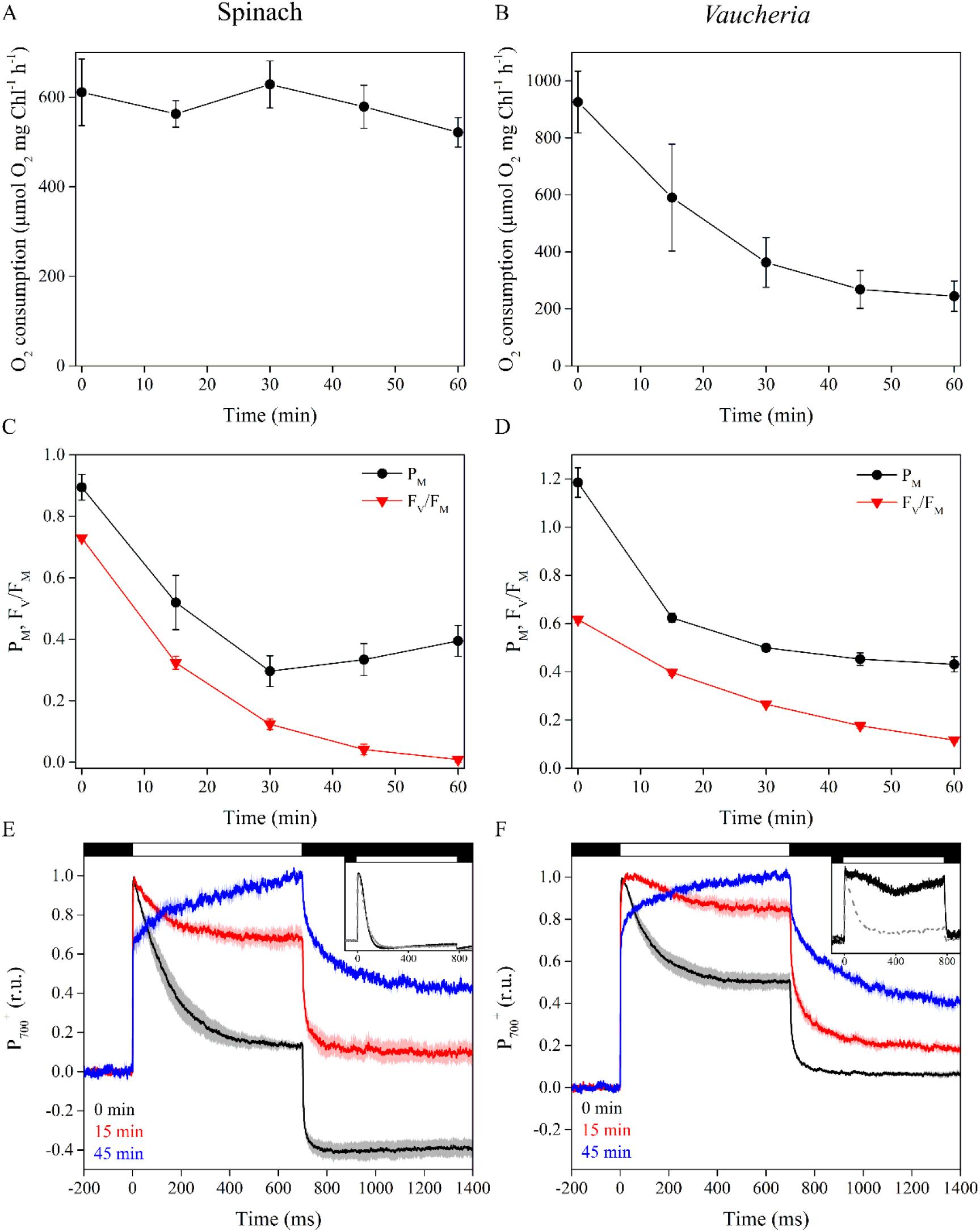
Photoinhibition of PSI in isolated thylakoids of spinach and *V. litorea* during high-light treatment, estimated with oxygen measurements or absorption based methods. (A, B) Photoinhibition of PSI in spinach and *V. litorea*, respectively, in the same experimental setup as in Fig. 5. (C, D) Decrease in maximal oxidation of P_700_ reaction center Chl (PM; black) and PSII photochemistry (F_V_/F_M_; red) during high-light treatment (PPFD 1000 μmol m^−2^s^−1^) in isolated spinach and *V. litorea* thylakoids. (E) P_700_ redox kinetics of spinach thylakoids during the high-light pulses used for P_M_ determination after 0, 15 and 45 min in photoinhibition treatment (black, red and blue, respectively). Black and white bars on top indicate darkness and illumination by the high-light pulse, respectively. (F) The same measurements as in (E) in *V. litorea* thylakoids. The insets of (E, F) show P_700_ redox kinetics from intact spinach leaves and *V. litorea* cells in aerobic (black, solid line) and anaerobic conditions (grey, dashed line). Dark control experiments are shown in Supplementary figure S3. The P_700_ kinetics in (E) and (F) have been normalized to stress the form of the curve. All data are averages from at least three biological replicates and error bars and the shaded areas around the curves indicate SE.

In both spinach and *V. litorea*, redox kinetics of P_700_, measured in aerobic conditions from thylakoids (Fig. 6E,F) were similar as their respective *in vivo* kinetics (Fig. 6E,F insets), i.e. P_700_ in *V. litorea* remained more oxidized during a light pulse than in spinach. Isolating thylakoids from *V. litorea* did, however, cause a decrease in P_700_ oxidation capacity. Unlike in spinach, P_700_ remains oxidized during a high-light pulse in intact *V. litorea* cells if oxygen is present, indicating that alternative electron sinks, such as flavodiiron proteins, function as efficient PSI electron acceptors *V. litorea* (Fig. 6E,F insets), probably protecting PSI against formation of ROS (Allahverdiyeva *et al*., 2015; Ilík *et al*., 2017; Shimakawa *et al*., 2019). In both species, P_700_ redox kinetics changed in the same way during the course of the high-light treatment of isolated thylakoids. The tendency of both species to maintain P_700_ oxidized throughout the high-light pulse in measurements done after 15 min treatment in high light is possibly due to decreasing electron donation caused by photoinhibition of PSII. At 45 min timepoint the damage to PSI is more severe, as indicated by a clear slowing down of P_700_ oxidation, which could be associated with problems in electron donation to downstream electron acceptors of PSI, such as ferredoxin (Fig. 6E,F).

## Discussion

### Upregulation of FtsH at the center of *V. litorea* plastid longevity

Previous studies have shown that the kleptoplasts stemming from *V. litorea* carry out *de novo* protein translation and are generally quite robust inside *E. chlorotica* (Green *et al*., 2000; Rumpho *et al*., 2001; Green *et al*., 2005). Our transcriptomic analysis of *V. litorea* plastids demonstrates active and regulated transcription of the plastome throughout the seven days of isolation we tested (Fig. 2), deepening our knowledge about the factors underpinning their native robustness. Considering gene orientation of the up- and downregulated genes suggests that e.g. *ftsH* and *psbB*, neighboring genes sharing the same orientation, do not constitute an operon (Fig. 2A).

Our results highlight the upregulation of *ftsH* and *tufA* during a period of several days after isolation of *V. litorea* plastids. Active transcription of these genes also occurs in the plastids of *E. timida* after a month of starvation (de Vries *et al*., 2013). FtsH protease is critical for the PSII repair cycle, where it is responsible for degradation of the D1 protein after pulling it out of the PSII reaction center. Recent findings in cyanobacteria, green algae and higher plants imply that FtsH is also important for quality control of a multitude of thylakoid membrane proteins and thylakoid membrane biogenesis (reviewed by Kato and Sakamoto, 2018). These findings may suggest that already the removal of the D1 protein from damaged PSII serves to protect from further photodamage and the production of ROS. The results of our photoinhibition experiments on the long-term retention slug *E. timida* may serve as a model of photoinhibition in other slugs, as they indicate that the kleptoplasts of *E. timida* possess a genetic toolkit capable of maintaining a PSII repair cycle (Fig. 2C).

We showed that the capacity of *V. litorea* plastids to recover from photoinhibition of PSII in the presence of CHI is nearly unaffected (Fig. 3B). While our CHI experiments on spinach need further exploration in terms of CHI effects, studies on the green alga *Chlamydomonas reinhardtii* (that also lacks *ftsH* in its plastome) have shown severe defects in PSII repair both during high-light and subsequent recovery when exposed to CHI (Fig. 3A, Wang *et al*., 2017). *C. reinhardtii* mutant lines have also been used to show that abundant FtsH offers protection from photoinhibition of PSII and enhances the recovery process (Wang *et al*., 2017). In *C. reinhardtii*, the FtsH hetero-oligomers responsible for D1 degradation are comprised of FtsH1 (A-type) and FtsH2 (B-type) (Malnoë *et al*., 2014). We probed the relative FtsH protein levels of *V. litorea* during the photoinhibition experiment using antibodies raised against *A. thaliana* A-(FtsH 1+5) and B-type FtsH (FtsH 2+8) in the absence and presence of CHI (Fig. 3D). At the end of the recovery period the CHI treated cells showed elevated levels of FtsH according to both tested antibodies. The elevated FtsH abundance did not enhance the recovery from photoinhibition of PSII in our experimental setup (Fig. 3B), but our results do point to a tendency of both, truly isolated (Fig. 2) and functionally isolated (Fig. 3) *V. litorea* plastids, to upregulate FtsH.

### Low ^1^O_2_ yield does not prevent photoinhibition of PSII, but can help maintain efficient repair processes in *V. litorea*

A green alga that is nearly immune to photoinhibition of PSII, *Chlorella ohadii,* has been isolated from the desert crusts of Israel (Treves *et al*., 2013; 2016). Its resilience against photoinhibition of PSII has largely been attributed to very narrow energetic gap between Q_A_ and Q_B_, favoring non-radiative charge recombination pathways within PSII that do not lead to ^1^O_2_ production (Treves *et al*., 2016). While *V. litorea* does not have as small energetic gap between Q_A_ and Q_B_ as *C. ohadii* (temperature difference of *V. litorea* Q- and B-band thermoluminescence peaks was 10 °C, whereas in *C. ohadii* it is only 2-4 °C), PSII charge recombination reactions of *V. litorea* appear to be very slow compared to those of spinach (Fig. 5B-D). Furthermore, the low ^1^O_2_ yield in *V. litorea* (Fig. 5A) suggests that the charge recombination reactions favor the direct non-radiative pathway. The low ^1^O_2_ yield in *V. litorea* likely factors into the lower dark-corrected rate constant of PSII photoinhibition in comparison to that of spinach thylakoids (Table 2) (Vass, 2011). All of our experiments, however, show that *V. litorea* does experience quite regular levels of PSII photoinhibition. This could indicate that the most important effect of the low ^1^O_2_ yield is protection of the autonomous maintenance machinery of the plastids, as ^1^O_2_ has been shown to be specifically harmful for the PSII repair cycle (Nishiyama *et al*., 2004).

### *V. litorea* thylakoids are highly vulnerable to ROS in the absence of regular stromal electron sinks

Despite the lower rate constant of PSII photoinhibition (Table 2) and ^1^O_2_ yield (Fig. 5A), *V. litorea* thylakoids exhibited drastic oxidative damage to lipids and proteins under high light (Fig. 4C,D). Isolated thylakoids are stripped of the main electron sink of PSI, the Calvin-Benson-Bassham cycle, and comparing P_700_ redox kinetics of *V. litorea* cells and isolated thylakoids (Fig. 6F and inset) reveals that they are also, at least partially, devoid of a Mehler-like reaction that safely reduces oxygen to water (Allahverdiyeva *et al*., 2013). This suggests that catalysts of oxygen reduction in *V. litorea* are likely soluble and therefore lost during the isolation procedure. Angiosperm plants like spinach do not rely on a Mehler-like reaction and are susceptible to photoinhibition of PSI in fluctuating light (Shimakawa *et al*., 2019). The PSI photoprotection by Mehler-like reaction has been assigned to enhanced electron sink capacity that lowers the probability of one-electron reduction of oxygen to superoxide by PSI. In comparison to spinach, this would make intact plastids of *V. litorea* less reliant on other ROS detoxification components that detoxify superoxide and hydrogen peroxide in the water-water cycle (Asada, 1999). Conversely, loss of the Mehler-like reaction during thylakoid isolation would leave the thylakoids highly conducive for ROS production by PSI and very susceptible to oxidative damage of the entire photosynthetic machinery. This is likely behind the finding that *V. litorea* thylakoids lose the ability to reduce methyl viologen in a high-light treatment that does not affect spinach thylakoids (Fig. 6A,B). When damage to PSI was estimated as a decrease in PM, spinach and *V. litorea* thylakoids showed very similar responses to high light, with both species exhibiting a decrease in PSI activity until electron donation from PSII was diminished due to photoinhibition of PSII (Fig. 6C,D), as suggested earlier (Sonoike, 1995; 1996). This, in addition to the highly similar changes in the redox kinetics of P_700_ during the photoinhibition treatment (Fig. 6E,F) between the two species, would suggest that the decrease in oxygen consumption in *V. litorea* thylakoids is caused by a further, more severe damage to PSI than the process causing the decrease in PM. The nature of this reaction is not known but it may be caused by production of ROS due to continuing electron flow through PSI in thylakoids of *V. litorea* exhibiting a low rate constant of PSII photoinhibition (Table 2) and normally relying on stromal electron acceptors for protection of PSI.

PSI of *V. litorea* is not particularly prone to photoinhibition, but our results do confirm that the electron sinks of photosynthesis must be functional in order to avoid large scale oxidative damage. This is especially relevant for animals that host a foreign organelle where uncontrolled ROS production is detrimental (de Vries *et al*. 2015). Our recent results on the LTR slug *E. timida* show that oxygen functions as an alternative electron sink in the slug plastids (Havurinne and Tyystjärvi, 2020), but whether the record-holding *E. chlorotica* utilizes the oxygen dependent electron sinks provided by *V. litorea* (Fig. 6F inset) remains to be tested. As for the main electron sink of photosynthesis, the carbon fixation rates of the plastids inside *E. chlorotica* are comparable to the rates measured from *V. litorea* cells after incorporation (Rumpho *et al*., 2001), suggesting that carbon fixation is not a problem in *E. chlorotica*.

## Conclusion

Plastids of *V. litorea* are genetically more autonomous than those of embryophytes, containing genes that help to maintain plastid functionality. Isolating the plastids triggers upregulation of the translation elongation factor EF-Tu and the central maintenance protease FtsH – a phenomenon that may be important for plastid longevity in the foreign cytosol of a sea slug. Low ^1^O_2_ yield protects the functionality of the plastid-encoded maintenance machinery and may slow down photoinhibition of PSII. Interruption of oxygen dependent alternative electron sinks upstream of PSI leads to large scale oxidative damage in *V. litorea*, suggesting that carbon fixation, the main electron sink of photosynthesis, needs to remain in near perfect working order to avoid destruction of the plastids. Our results support decades old data (Trench *et al*., 1973 *a*, b) suggesting that the native stability and associated peculiar functionality of the plastids themselves hold the key to long-term kleptoplast longevity in sacoglossans. Nature has evolved an elaborate suite of photoprotective mechanisms and the unique animal-kleptoplast association allows to explore them and even identify new ones.

## Abbreviations

^1^O_2_: singlet oxygen
CHI: cycloheximide
DCBQ: 2,6-dichloro-1,4-benzoquinone
DCMU: 3-(3,4-dichlorophenyl)-1,1-dimethylurea
DCPIP: 2,6-dichlorophenolindophenol
F_V_/F_M_: maximum quantum yield of PSII photochemistry
k_PI_: rate constant of PSII photoinhibition
MDA: malondialdehyde
OEC: oxygen evolving complex of PSII
P_680_: reaction center Chl of PSII
P_700_: reaction center Chl of PSI
P_M_: maximum oxidation of P700
PPFD: photosynthetic photon flux density
PSI: Photosystem I
PSII: Photosystem II
ROS: reactive oxygen species
TEM: transmission electron microscope
TyrD^+^: oxidized tyrosine-D residue of PSII

## Supplementary data

Supplementary data are available at *JXB* online.

*Table S1.* List of primers used in qPCR experiment.

*Table S2.* Modeled S-state distribution of the OEC in spinach and *V. litorea*.

*Fig. S1.* EPR spectra from spinach and *V. litorea* thylakoids.

*Fig. S2.* Dark control treatments of *in vitro* PSII photoinhibition in spinach and *V. litorea*.

*Fig. S3.* Dark control treatments of *in vitro* PSI and PSII photoinhibition in spinach and *V. litorea*.

*Fig. S4. In vitro* PSI and PSII photoinhibition in DCMU treated spinach thylakoids.

## Acknowledgements

This work was supported by Academy of Finland (grant 333421, ET). VH was supported by Finnish Cultural Foundation, Finnish Academy of Science and Letters (Vilho, Yrjö and Kalle Väisälä fund), Turku University Foundation and University of Turku Graduate School. Sofia Vesterkvist is thanked for help with *E. timida* photoinhibition measurements. SBG would like to thank the German research council for funding through the CRC 1208-267205415 – and the SPP2237, and the group of U.-G. Maier (Marburg) for help with electron microscopy.

## Author Contributions

VH, SBG and ET planned the experiments. VH did all photosynthesis and ^1^O_2_ measurements and wrote the paper with comments from all authors; MH, supervised by SBG, did the gene expression measurements and TEM imaging; MA measured lipid peroxidation, protein oxidation and EPR spectra; SK developed the cuvette system for P_700_^+^ measurements; ET supervised the work.

## Data Availability

The data that support the findings of this study are openly available in Mendeley Data at http://doi.org/10.17632/535dcxjt2d.1.

---

**Supplementary table S1.**
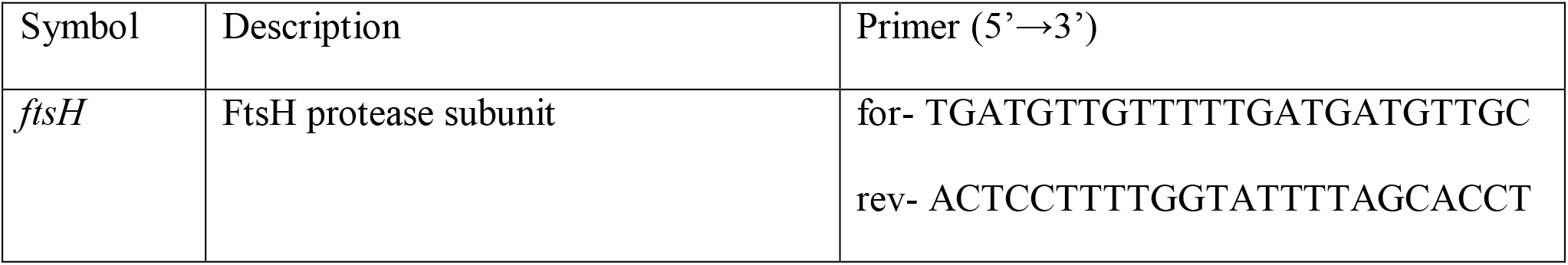
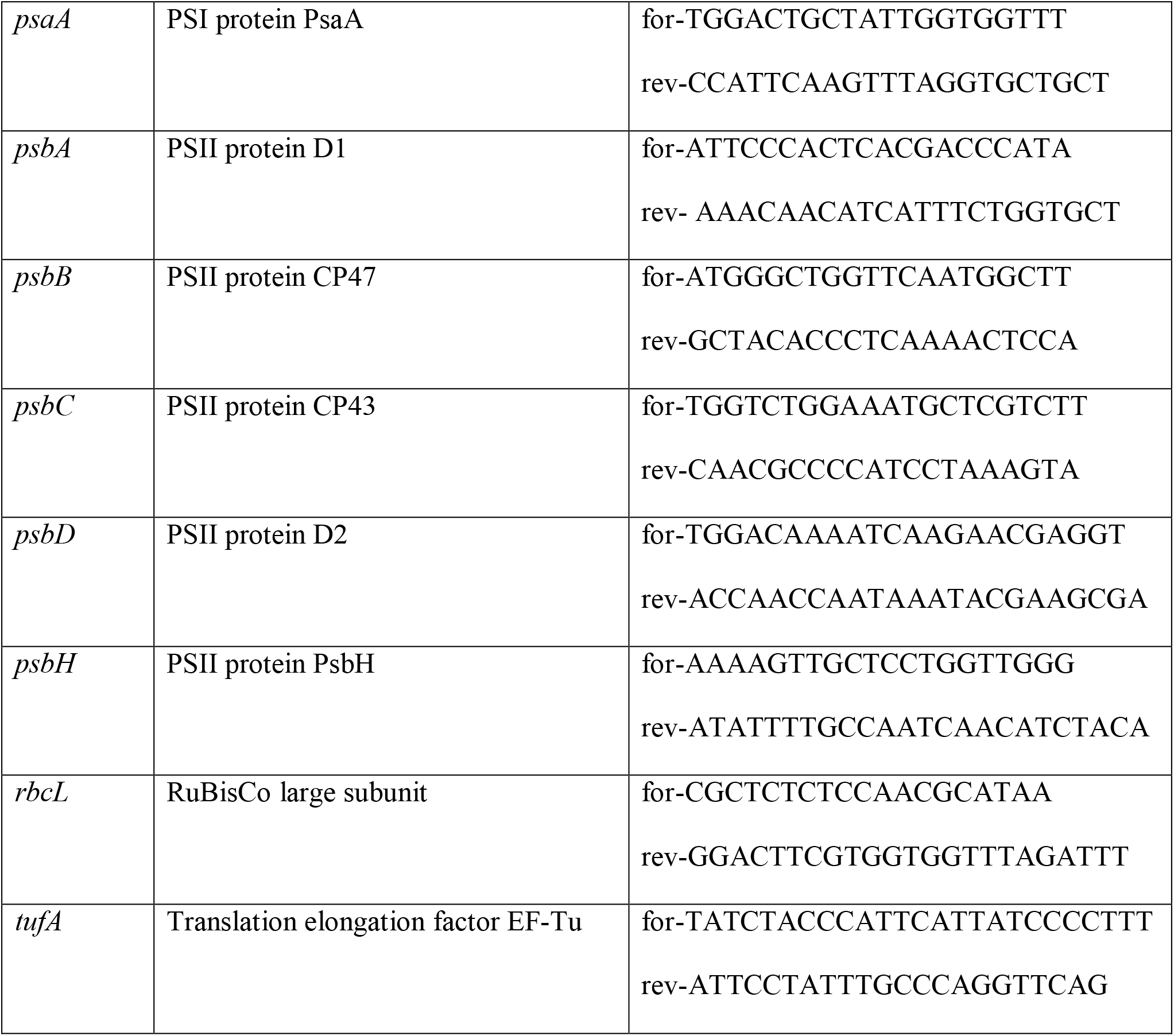
The list of primers designed for quantitative real-time PCR analysis of transcription in isolated *V. litorea* plastids.

**Supplementary table S2.** Percentage distribution of the S-states of the OEC in isolated thylakoids from spinach and *V. litorea* after different preflash treatments prior to measuring flash induced oxygen evolution. The flash oxygen data in Figure 5B was modeled essentially as described in Antal *et al*., (2009) to estimate the S-state distribution.

|  | S<sub>0</sub> % (±SE) | S<sub>1</sub> % (±SE) | S<sub>2</sub> % (±SE) | S<sub>3</sub> % (±SE) |
| --- | --- | --- | --- | --- |
| Spinach 10 min dark | 0.01 (±0.01) | 71.01 (±2.78) | 24.54 (±1.95) | 4.44 % (±1.15) |
| Spinach preflash + 10 s dark | 0 | 32.30 (±9.36) | 57.28 (±5.83) | 10.42 (±3.63) |
| Spinach preflash + 100 s dark | 0 | 65.30 (±0.17) | 26.32 (±0.07) | 8.39 (±0.24) |
| *Vaucheria* 10 min dark | 6.84 (±3.20) | 63.44 (±3.66) | 24.65 (±3.70) | 5.08 (±3.20) |
| *Vaucheria* preflash + 10 s dark | 0.02 (±0.01) | 24.69 (±1.19) | 60.62 (±0.59) | 14.66(±0.83) |
| *Vaucheria* preflash + 100 s dark | 0.01 (±0.01) | 45.21 (±2.03) | 39.18 (±0.92) | 15.61 (±2.53) |

**Supplementary Figure S1.**
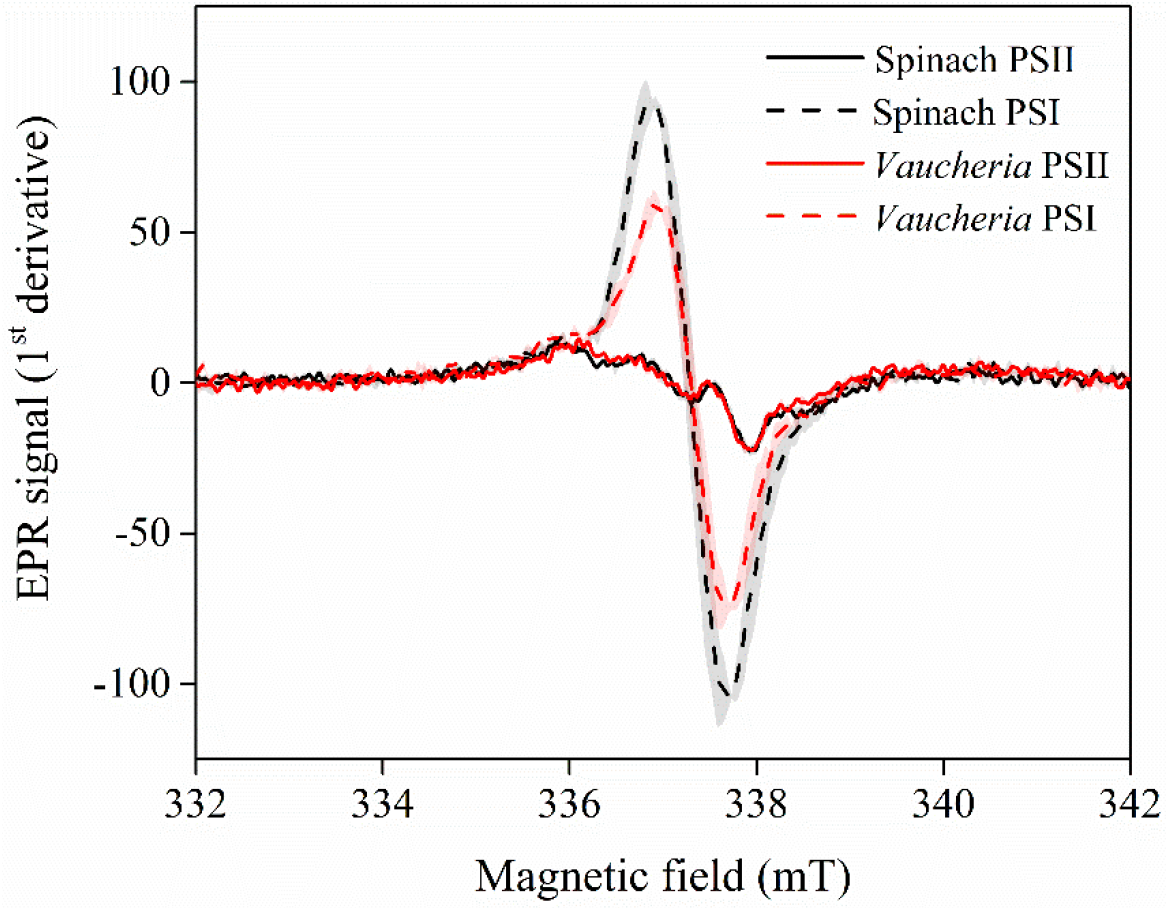
EPR spectra of PSII (Tyr_D_^+^) and PSI (P_700_^+^) in spinach and *V. litorea* thylakoids. All spectra were measured from isolated thylakoid samples containing 2000 μg total Chl ml^−1^. Each curve represents an average of three independent biological replicates and the shaded areas around the curves represent SE.

**Supplementary Figure S2.**
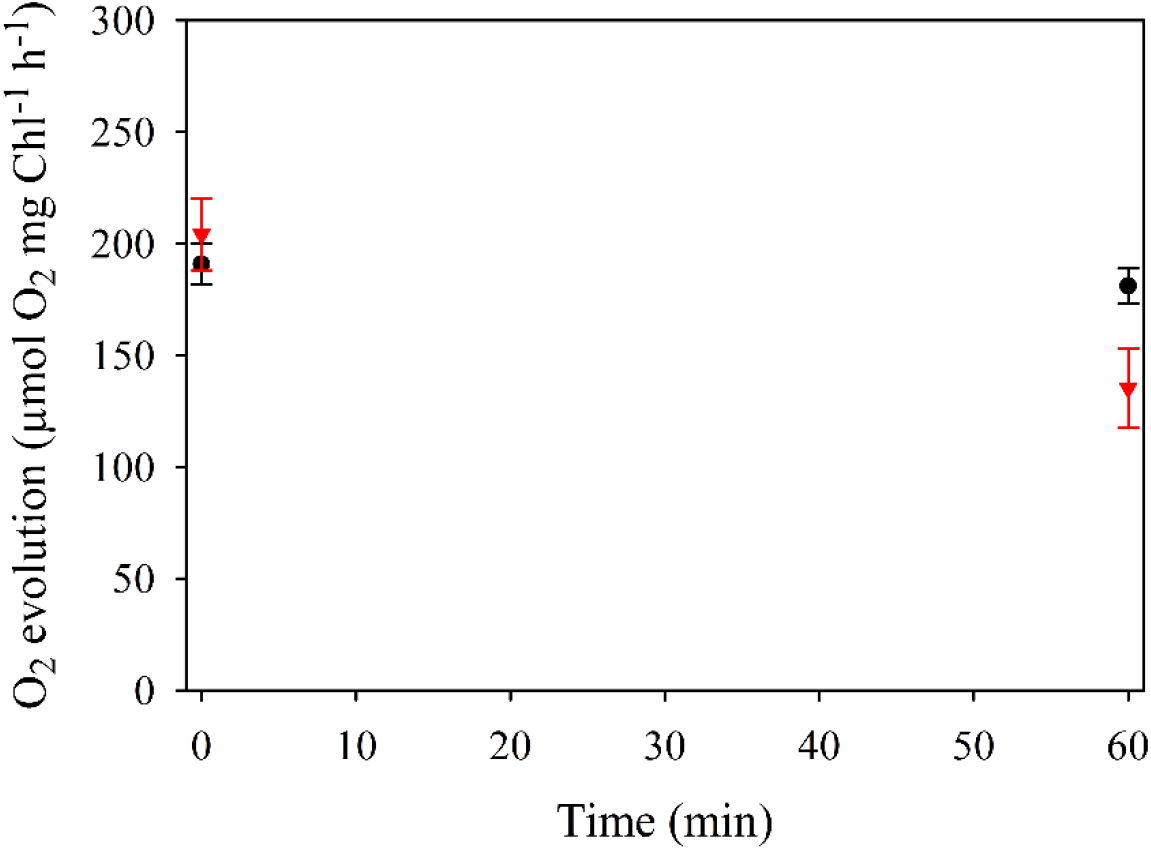
Dark control treatments of the *in vitro* photoinhibition experiments shown in Fig. 4A of the main text. PSII activities of spinach (black) and *V. litorea* (red) at the onset and after a 60 min dark treatment at 22 °C in photoinhibition buffer. Oxygen evolution was measured in the presence of 0.5 mM DCBQ and hexacyanoferrate(III) from samples containing 20 μg total Chl ml^−1^. Rate constant of PSII dark inactivation was 0.001 min^−1^ for spinach and 0.007 min^−1^ for *V. litorea*. Each data point represents an average of three biological replicates and the error bars indicate SE.

**Supplementary figure S3.**
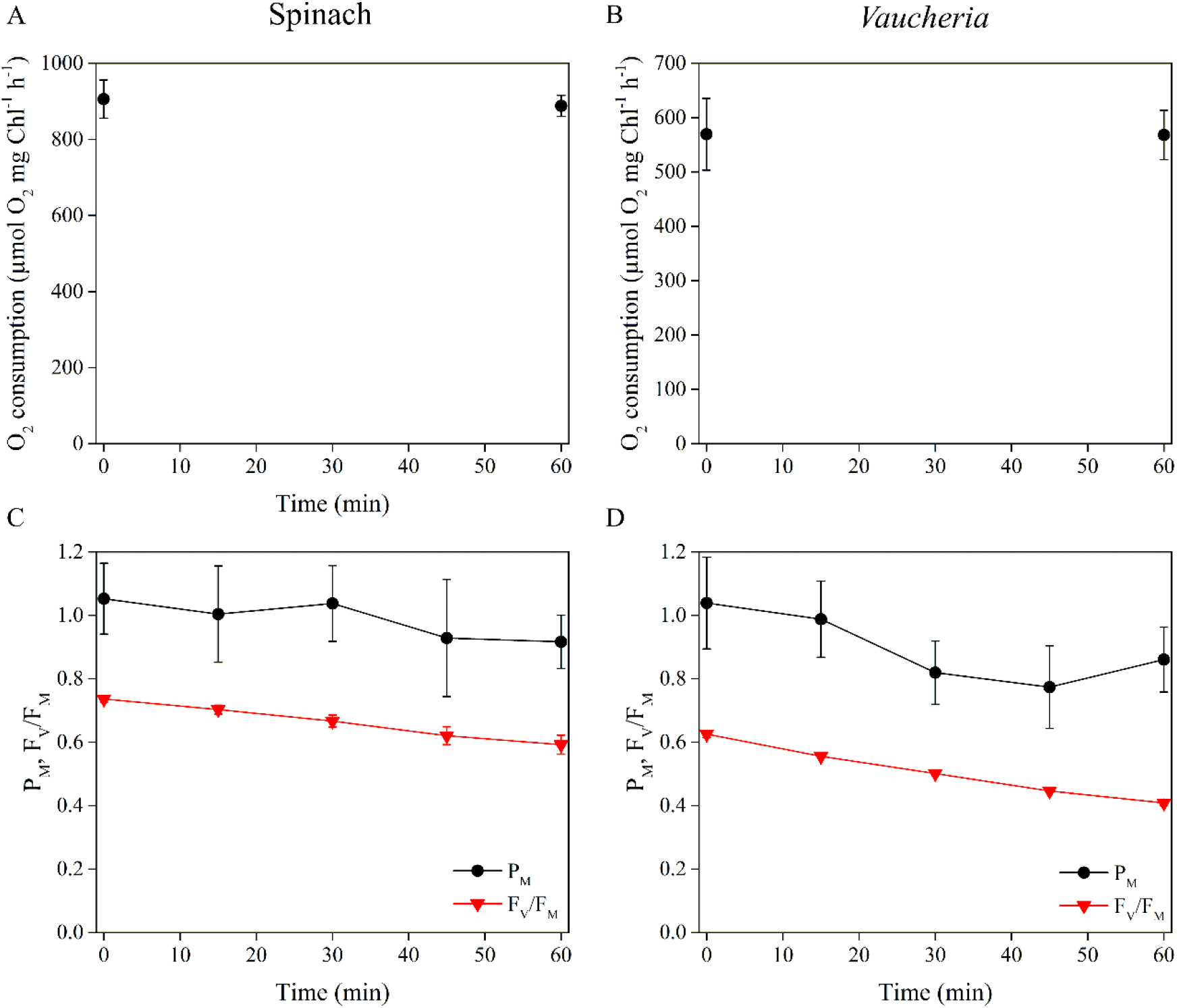
Dark control treatments of the *in vitro* photoinhibition experiments shown in Fig. 6 of the main text. (A, B) PSI activities of spinach and *V. litorea* during a 60 min dark incubation period in photoinhibition buffer, measured as oxygen consumption. (C, D) PSI and PSII activities in isolated spinach and *V. litorea* thylakoids during a 60 min dark treatment, as estimated by maximal oxidation of P_700_ (PM; black) and F_V_/F_M_ (red), respectively. All data are averages from a minimum of three biological replicates and error bars indicate SE.

**Supplementary figure S4.**
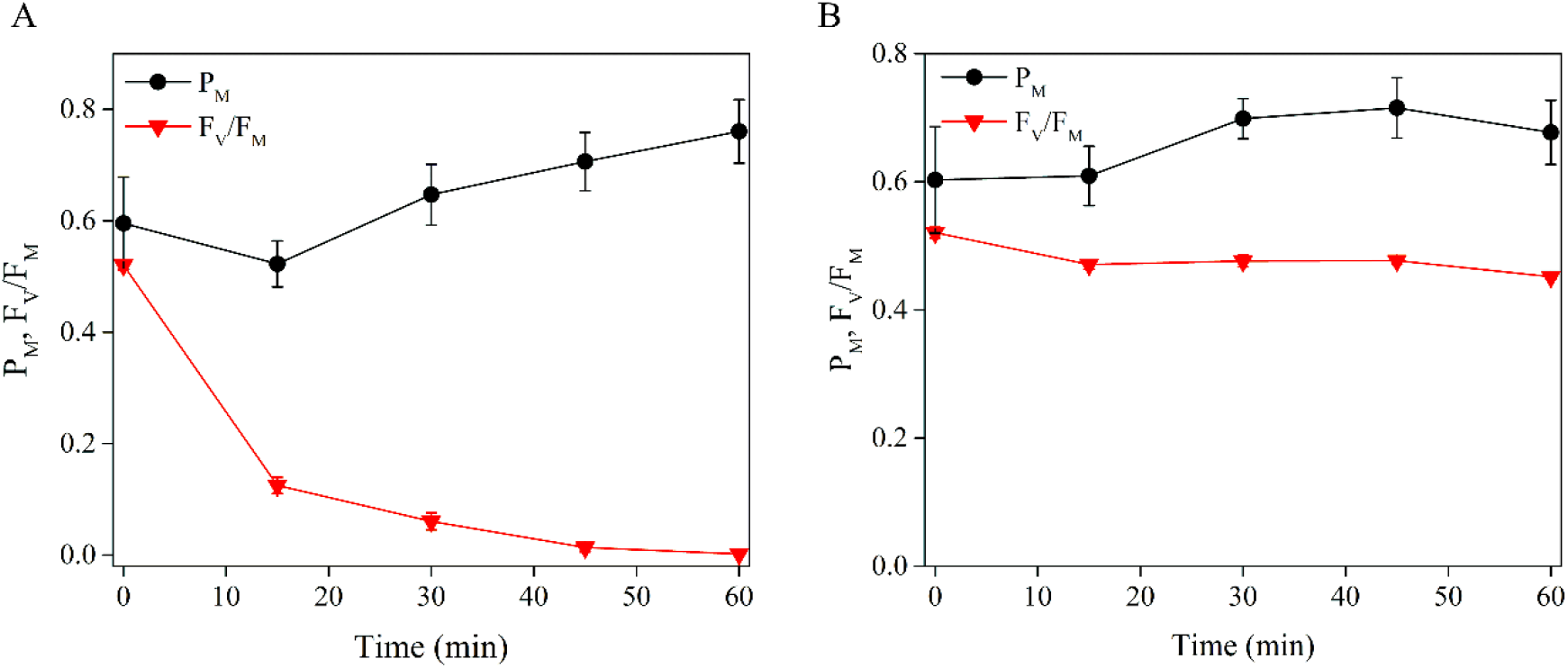
DCMU prevents photoinhibition of PSI in isolated spinach thylakoids. (A) Maximal oxidation of P_700_ (PM; black) and maximum quantum yield of PSII (F_V_/F_M_; red) were measured during a 60 min high-light treatment (PPFD 1000 μmol m^−2^s^−1^) in DCMU treated spinach thylakoids. (B) Dark control experiments using the same setup as in (A). All data are averages from four biological replicates and the error bars indicate SE.

